# PGCN: Disease gene prioritization by disease and gene embedding through graph convolutional neural networks

**DOI:** 10.1101/532226

**Authors:** Yu Li, Hiroyuki Kuwahara, Peng Yang, Le Song, Xin Gao

**Author notes:** All correspondense should be addressed to Xin Gao and Le Song.

## Abstract

**Motivation:** Proper prioritization of candidate genes is essential to the genome-based diagnostics of a range of genetic diseases. However, it is a highly challenging task involving limited and noisy knowledge of genes, diseases and their associations. While a number of computational methods have been developed for the disease gene prioritization task, their performance is largely limited by manually crafted features, network topology, or pre-defined rules of data fusion.

**Results:** Here, we propose a novel graph convolutional network-based disease gene prioritization method, PGCN, through the systematic embedding of the heterogeneous network made by genes and diseases, as well as their individual features. The embedding learning model and the association prediction model are trained together in an end-to-end manner. We compared PGCN with five state-of-the-art methods on the Online Mendelian Inheritance in Man (OMIM) dataset for tasks to recover missing associations and discover associations between novel genes and diseases. Results show significant improvements of PGCN over the existing methods. We further demonstrate that our embedding has biological meaning and can capture functional groups of genes.

**Availability:** The main program and the data are available at https://github.com/lykaust15/Disease_gene_prioritization_GCN.

## 1 INTRODUCTION

The last decade has seen a rapid increase in the adoption of whole-exome sequencing in the clinical diagnosis of genetic diseases (Feero, 2014). However, the success rate of such genome-based diagnostics still remains far from perfect, with reported yields for a range of Mendelian diseases ranging from ∼20 to ∼50% (Taylor *et al.*, 2015; Retterer *et al.*, 2016). This relatively low yield is largely attributed to a considerable difficulty in differentiating disease-causing variants from a large pool of rare genetic variants that are not pathogenic and do not play roles in the expression of the disease phenotype (MacArthur *et al.*, 2014; Tranchevent *et al.*, 2016). To efficiently detect pathogenic variants and to improve the diagnostic rate of the genome-based approach, it is essential to have disease gene prioritization that substantially reduces the number of candidate causal variants and ranks them for further interrogations based on the association of the corresponding genes with the disease phenotype.

A number of computational methods have been developed to tackle the disease gene prioritization problem (Wang *et al.*, 2011; Moreau and Tranchevent, 2012), and have been shown to be useful. For example, Endeavour (Aerts *et al.*, 2006; Tranchevent *et al.*, 2008, 2016) was able to associate *GATA4* with congenital diaphragmatic hernia (Yu *et al.*, 2013); GeneDistiller (Seelow *et al.*, 2008) discovered the role of *MED17* mutations in infantile cerebral and cerebellar atrophy (Kaufmann *et al.*, 2010). Based on the underlying computational techniques, existing disease gene prioritization methods can be categorized into five types. The first type is filter methods (Franke *et al.*, 2004; Bush *et al.*, 2009; Mordelet and Vert, 2011; Deo *et al.*, 2014), which sift the candidate list of genes into a smaller one according to the properties that associated genes should have. The second type of methods is based on text mining (Perez-Iratxeta *et al.*, 2005; Yu *et al.*, 2008; ElShal *et al.*, 2016; Smaili *et al.*, 2018a,b). Such methods score the candidate genes using the co-occurrence evidence with a certain disease from the literature. Thus, these methods can only detect associations that are already known. The third type is similarity profiling and data fusion methods (Aerts *et al.*, 2006; De Bie *et al.*, 2007; Chen *et al.*, 2009; Gefen *et al.*, 2010; Li and Patra, 2010; Britto *et al.*, 2012; Zitnik *et al.*, 2015; Zakeri *et al.*, 2015; Kim *et al.*, 2015; Tranchevent *et al.*, 2016; Kumar *et al.*, 2018). This is the dominant type in the disease gene prioritization community and includes the famous Endeavour method (Aerts *et al.*, 2006). These methods are based on the idea that similar genes should be associated with similar sets of diseases and vise versa. The similarity measurement can be defined using different data sources, such as Gene Ontology (GO) or the BLAST score. After obtaining the similarity scores from each data source, such methods apply data fusion to aggregate these scores into a global ranking. The fourth type is network-based methods (Wang *et al.*, 2011; Lee *et al.*, 2011; Guan *et al.*, 2012; Li and Li, 2012; Magger *et al.*, 2012; Kacprowski *et al.*, 2013; Nitsch *et al.*, 2011; Singh-Blom *et al.*, 2013; Rao *et al.*, 2018). Such methods represent diseases and genes as nodes in a heterogeneous network, in which the edge weight represents their similarities. The last type is based on matrix completion techniques in recommender systems (Natarajan and Dhillon, 2014; Zakeri *et al.*, 2018). These methods represent the disease gene association as an incomplete matrix and solve the disease gene prioritization problem by filling the missing values of the matrix. This category of methods has been shown to be the state-of-the-art (Zakeri *et al.*, 2018).

Despite the advances of the existing efforts, they have the following bottlenecks. Firstly, the similarity-based methods, which are rooted in the “guilt-by-association” principle, often fail to handle new diseases whose associated genes are completely unknown (Zakeri *et al.*, 2018). Secondly, although the performance of the network-based methods is reasonable, they are biased by the network topology and cannot easily integrate multiple sources of information about genes and diseases (Moreau and Tranchevent, 2012). Thirdly, matrix completion methods assume and look for a weighted linear relationship between genes and diseases, which, in reality, is most likely to be highly nonlinear (Navlakha and Kingsford, 2010). In addition, most of the existing methods rely heavily on manually-crafted features or pre-defined rules of data fusion. Therefore, the disease gene prioritization problem remains elusive. On the other hand, the recent success of graphical models and deep learning in bioinformatics (Zitnik *et al.*, 2018; Li *et al.*, 2017; Dai *et al.*, 2017; Kim *et al.*, 2018; Xia *et al.*, 2018) suggests the possibility to systematically incorporate multiple sources of information in the heterogeneous network and learn the highly nonlinear relationship between diseases and genes.

In this paper, we propose a novel disease gene prioritization method, PGCN, based on graph convolutional neural networks (GCN) (Dai *et al.*, 2016; Kipf and Welling, 2016; Hamilton *et al.*, 2017; Zitnik *et al.*, 2018). Starting from a heterogeneous network which is composed of the genetic interaction network, the human disease similarity network, and the known disease-gene association network, with the additional information about genes and diseases from multiple sources, our method first learns embeddings for genes and diseases through graph convolutional neural networks, by considering both the network topology and the additional information of diseases and genes. Such embeddings are fed into an edge decoding (edge prediction) model to make predictions for disease gene associations. Although we describe our method in two steps, our model is trained in an end-to-end manner so that the model can learn the embedding and the decoding jointly.

We compared PGCN with five state-of-the-art methods (GeneHound (Zakeri *et al.*, 2018), IMC (Natarajan and Dhillon, 2014), GCAS (Rao *et al.*, 2018), Catapult (Singh-Blom *et al.*, 2013), and Katz (Singh-Blom *et al.*, 2013)) on the Online Mendelian Inheritance in Man (OMIM) dataset (Hamosh *et al.*, 2005). Extensive experiments suggest that our method significantly outperforms the existing methods on recovering missing associations, and on discovering associations for novel genes and/or diseases that are not seen in the training. We further demonstrate that our embedding has biological meaning and can capture functional groups of genes.

## 2 METHODS

In our work, we cast the disease gene prioritization problem as a link prediction problem. Unlike the previous studies (Natarajan and Dhillon, 2014; Zakeri *et al.*, 2018) which solve the problem with matrix factorization, we propose a novel method based on graph convolutional neural networks. We compile the disease similarities, genetic interactions, and disease-gene associations into a multi-nodal heterogeneous network, as shown in Fig. 1. In this network, the potential disease-gene associations can be considered as missing links and our goal is to predict these links (Chen *et al.*, 2005; Ying *et al.*, 2018). The overview of our method is shown in Fig. 2. The core idea of our method is to learn the nodes’ latent representations (embeddings) from their initial raw representations (information encoded from different sources), considering the graph’s topological structure and the nodes’ neighborhood, after which we make predictions using the learned embeddings with the edge decoding model. Both the embedding model and the decoding model are trained in an end-to-end manner so that each model is optimized while being regularized by the other one. In the following sections, we introduce each component of the proposed method in more details.

**Fig. 1:**
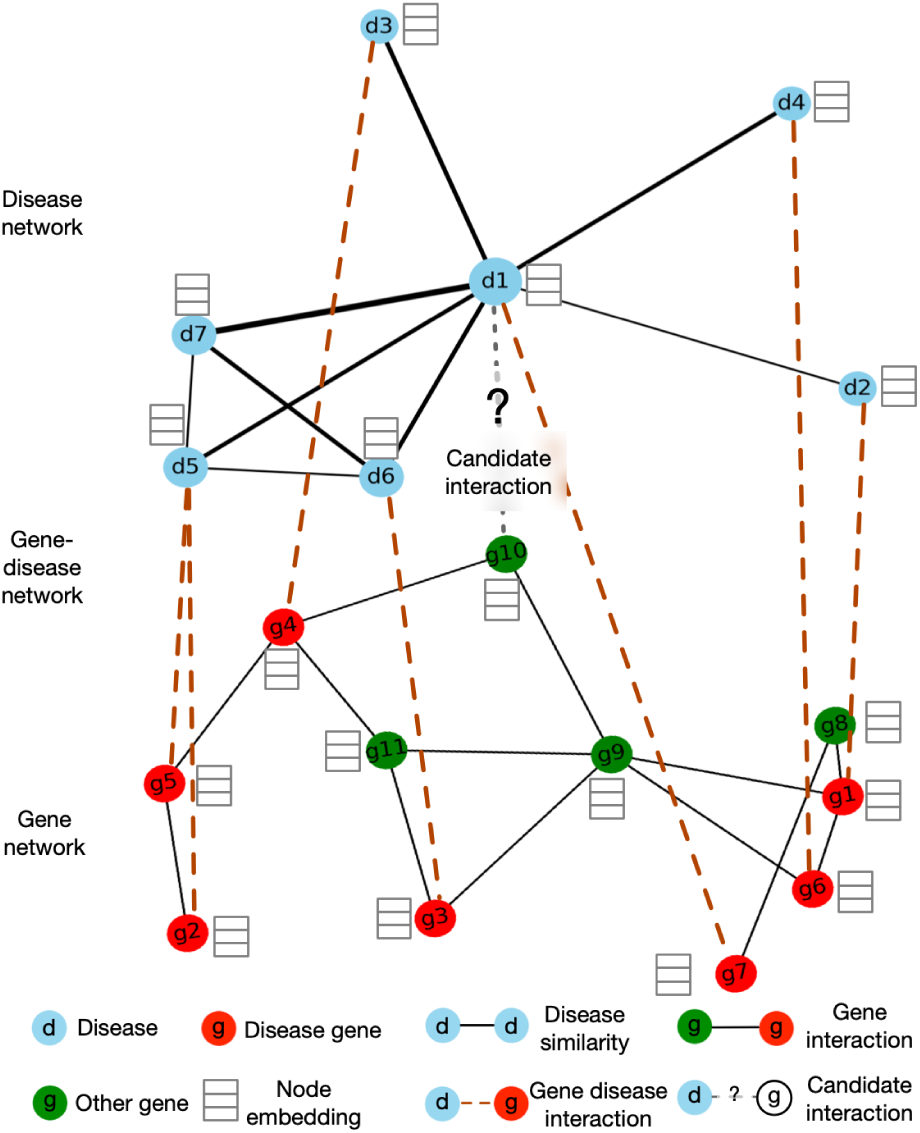
Disease gene prioritization as a link prediction problem. The heterogeneous network contains three components, the genetic interaction network, the disease similarity network, and the disease-gene association network. The potential disease gene associations can be considered as missing links in the disease-gene association network. Our goal is to predict those links given the heterogeneous network and additional raw representations of the nodes (diseases and genes).

**Fig. 2:**
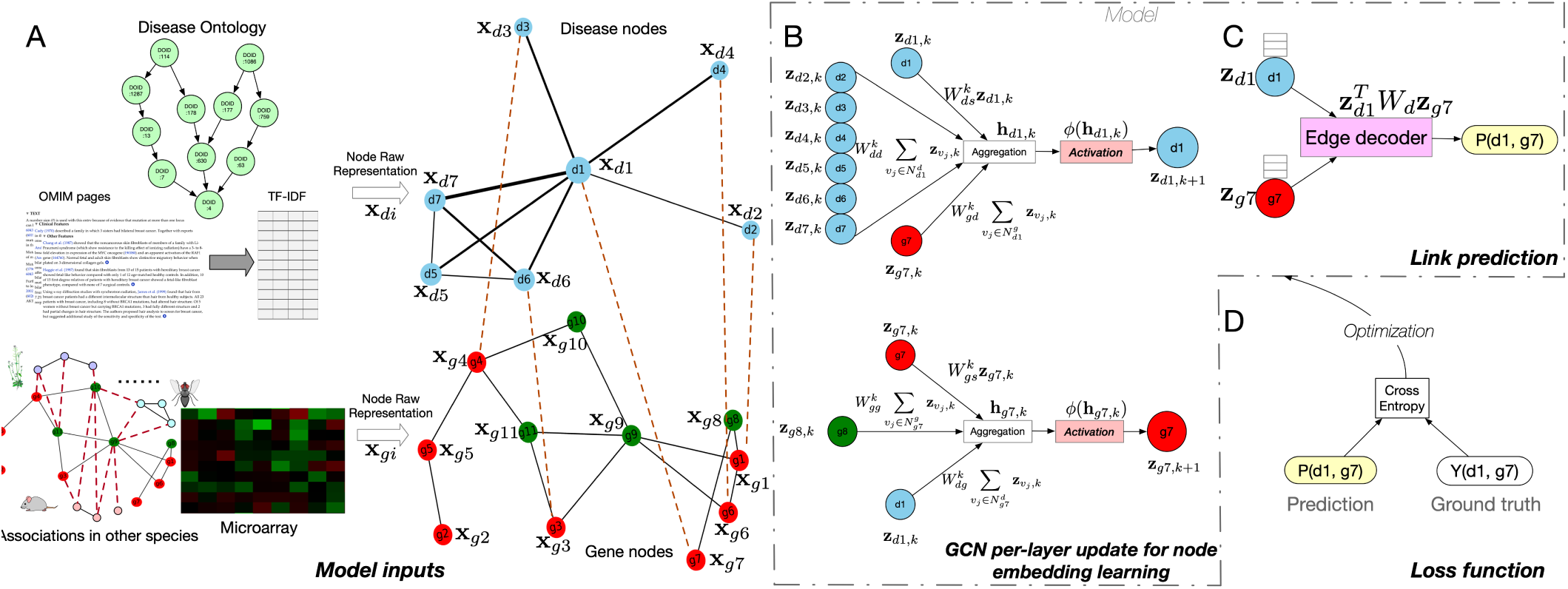
Overview of the proposed method. (A) The input of our model contains two components, the heterogeneous network and the additional information for the nodes. As for the heterogeneous network, we used HumanNet as the gene network, disease similarity network as the disease network, and the associations from OMIM as the disease-gene network. For the additional information of diseases, we used Disease Ontology similarity and the TF-IDF calculated from OMIM. For the additional information of genes, we used association matrices from other species and the gene expression microarray data. (B) Examples of one layer of the graph convolutional neural network update for learning node embeddings. For each node, the model aggregates information from its neighbor nodes’ previous layer embeddings and then apply activation to obtain the current layer embedding of that node. Note that for different nodes, the computational graphs can be different but the parameters are shared for the same operation in different computational graphs. (C) The link prediction model. We model the edge prediction from the learned node embeddings with bilinear edge decoder. (D) The cross-entropy loss calculated from the ground truth and the output of the link prediction model for certain edges (or non-edges) is used as the loss function to train both the node embedding model and the edge decoding model jointly in an end-to-end fashion.

### 2.1 Disease gene prioritization as a link prediction problem

Recent studies (Natarajan and Dhillon, 2014; Zakeri *et al.*, 2018) have formulated the disease gene prioritization problem as a matrix completion problem and applied the recently developed methods in recommender systems, resulting in better performance than the previous state-of-the-arts. Although we also consider the problem as a recommender system problem, we treat the entire data structure as a heterogeneous network (Fig. 1 and Section 2.2). Each node represents a disease or a gene, and each edge represents one specific kind of interaction. In addition, each disease and gene is supplemented with additional information from different data sources (Section 2.3). Our goal is to predict the potential links between disease nodes and gene nodes, whose link strength can be used for prioritization. Compared with the matrix factorization methods, our formulation can capture the nonlinear relationship between diseases and genes. Compared to the traditional network-based methods, our method is able to integrate the information from different sources in a systematic and natural way.

The core component of our method is the graph convolutional encoder (Section 2.4), which can learn the embeddings from the nodes’ neighborhood, node-specific information, and the topology of the heterogeneous network. The central problem for learning embeddings from graph data is to propagate and transform information. As shown in Fig. 2 (A), the entire graph starts from a heterogeneous network, with each node containing information from different sources. In the graph convolution model, each node’s neighboring nodes define the computational graph of its local neural network, i.e., its own neural network architecture. Although the local computational graphs can be different for different nodes, the same operations share the same parameters and activation functions, which specify how the information is shared and propagated across the computational graph. Since we instantiate the graph convolution operation using a fully-connected neural network (Fig. 2 (B)), the model can seamlessly integrate information from different sources. The embeddings are fed into the link decoding model (Section 2.5). Thus, the proposed method can achieve problem-specific data integration systematically, whose parameters are learned from the data in an end-to-end manner.

### 2.2 Network compiling

The network in our model (Fig. 1) is a heterogeneous network containing three components: the gene network, the disease similarity network, and the disease-gene network. The disease-gene network is built from the OMIM database (November 26, 2017), with the associations being the links. After preprocessing, this network contains 12331 genes, 3215 diseases, and 3988 disease-gene associations.

As for the gene network, we used HumanNet from Lee *et al.* (2011). This large-scale functional gene network was constructed by considering multiple sources of information, including human mRNA co-expression, protein-protein interactions, protein complex, and comparative genomics information. In total, it incorporated 21 genomics and proteomics datasets from four species. Compared to the network built from single dataset, such as protein-protein interaction networks, it has higher accuracy and genome coverage (Lee *et al.*, 2011). The usefulness of HumanNet in disease gene prioritization has been proved by previous studies (Singh-Blom *et al.*, 2013; Natarajan and Dhillon, 2014). In summary, our gene network is composed of 12331 genes and 733836 edges with positive weights. More details about the network can be found in Lee *et al.* (2011).

We used the MimMiner from Van Driel *et al.* (2006) as the disease similarity network. This network was built by using text mining analysis on the OMIM database. For each disease, the anatomy and disease sections of the medical subject headings were used to extract terms from OMIM, whose frequencies were used as the feature vectors of the disease. After further refinement, the feature vectors were used to compute the pairwise similarities between the disease, which resulted in the MimMiner network. Although in the construction process, it did not involve gene information, the similarities were shown to be positively correlated with a number of measures of gene function. This network has also been used as a feature input in the previous disease gene prioritization methods (Singh-Blom *et al.*, 2013; Natarajan and Dhillon, 2014). After setting the similarity threshold as 0.2, we obtained a disease similarity network with 3215 diseases and 645945 edges.

### 2.3 Data sources for node raw representation

In contrast to the other network-based methods, our model can naturally incorporate additional information about the nodes from different sources. In our implementation, we incorporated the following data sources, although our method is generic and can take any source of information for diseases and genes.

As shown in Fig. 2 (A), we incorporated two kinds of additional information for the disease nodes. The first data source is Disease Ontology (DO) similarity. After collecting the ontology for the disease nodes, we calculated a similarity matrix for those diseases using the Resnik pairwise similarity (Resnik, 1995) with the best-match average (BMA) strategy (Wang *et al.*, 2007). For each disease, we took the corresponding row of this matrix as an additional feature vector for this node. The second data source is the clinical text from OMIM webpages. We collected the Clinical Feature and Clinical Management sections from the OMIM webpages for each disease, and we removed the most frequent and most rare words. Then, we counted the frequency of each unique word in the corpus related to each disease. To remove the bias of the relatively frequent words, we applied the TF-IDF scheme to the term frequency matrix and obtained the corresponding row as the feature vector for a disease. Finally, the two vectors were concatenated as the additional information for the disease.

We also used two kinds of features as the additional information for the gene nodes. Following the strategy from Natarajan and Dhillon (2014), we collected the microarray measurement of gene expression level in different tissue samples from BioGPS and Connectivity Map. Since some genes are missing in the probes, we obtained 4536 features for 8755 genes. It is well-known that samples from the same cell type of different individuals tend to have a similar expression pattern, which results in redundant information in the obtained feature matrix. To eliminate the redundancy and reduce the dimensionality, we applied principle component analysis (PCA) on the features and used the first 100 eigenvectors as the feature representations from gene expression microarray. The second type of additional information for genes is derived from gene-phenotype associations of other species. Following the previous studies (Singh-Blom *et al.*, 2013; Natarajan and Dhillon, 2014), we used the phenotypes from eight species. As a result, we obtained eight matrices, whose rows represent different genes and columns represent the phenotypes of different species. We concatenated those gene-phenotype matrices together with the microarray matrix along the gene dimension, resulting in the additional information of the genes.

### 2.4 Node embedding with graph convolution

In this section, we introduce how we obtain the embeddings using graph convolutional neural networks, taking into consideration both the network topology, nodes’ neighborhood, and the additional information of the nodes. Formally, given a graph 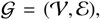, where 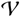 represents the set of nodes and 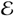 represents the set of edges, with the adjacent matrix as **A**, we denote **x**_*i*_ ∈ ℝ^*m*_*i*_^ as the additional information of the node 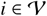. Note that in our method, the value of *m*_*i*_, which represents the dimension of the additional feature vectors can be different for different kinds of nodes, i.e., gene nodes and disease nodes. The goal of embedding is to map each node to a vector **z**_*i*_ ∈ ℝ^*c*^, where *c* ≪ *m*_*i*_, considering the information contained in **A** and 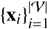.

The central problem of learning embedding with graph convolutional neural networks is to learn how to transform and propagate information (the additional information and intermediate embeddings of each node) across the entire network. In our method, the GCN module defines the information propagation architecture (the local computational graph) for each node using the node’s neighborhood in the graph 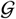. In terms of the parameterization of the local computational graph, which defines how the information is propagated and shared, the parameters and weights are shared across all the local computational graphs built from 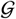, with the assumption that within the same graph 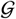, the way of sharing and propagating information should be the same. As a result, for a given node, each layer of graph convolutional neural networks aggregates and transforms the information (feature representations) from its neighbors and applies the same transformation to all part of the network, which is illustrated in Fig. 2 (B). If there is only one layer of graph convolution, the embedding will only aggregate information from its first-order neighbors. Thus, stacking *N* layers of graph convolutional layers can make the embedding effectively convolve information from its *N*-order neighbors explicitly. Besides, when we stack more than one graph convolutional layers, the information of each single node can start broadcasting to the entire network implicitly, whose affect depends on the network topological structure (size, connectivity *etc.*). By using multiple convolutional layers, we are able to learn the embedding of nodes, considering the network topology, local neighborhoods, and additional information of the nodes.

Formally, in each layer, for each node, the information aggregation and transformation model takes the following form:

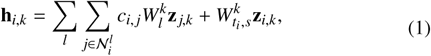

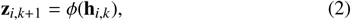

where **z**_*i*,*k*_ ∈ ℝ^*c*_*k*_^ is the hidden representation of node *i* in the *k*-th graph convolutional layer and *c*_*k*_ is the dimensionality of that hidden representation; **h**_*i*,*k*_ represents the feature vector which has aggregated the information from the *k*-th layer hidden representations of the node’s neighbors; *l* represents the link type, *i.e.*, genetic interaction, disease-disease similarity, or disease-gene association; 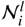 are the neighbors of *i*, which are linked by the link type *l*; 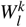 is the weight parameter related to the link type *l*, such as 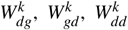 and 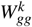 in Fig. 2 (B); *c*_*i,j*_ is the normalization constant, inspired by Zitnik *et al.* (2018), which is defined as 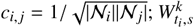 is the weight parameter preserving the information from the node itself, where *t_i_* indicates the type of the node; φ is the non-linear activation function, which is usually chosen as rectified linear unit (ReLU). Note that the above aggregation and transformation formulas are related to the neighbors of a certain node, which means that the computational graph architecture can be different for nodes with different local neighborhood structure. We show examples of two very different computational graphs for nodes *d*1 and *d*7 in Fig. 2 (B). On the other hand, although the computational graphs can be different, the parameters are only related to the link type, not related to the node neighborhoods, which means that the parameterization is shared across the entire graph.

In our method, we use summation as the information aggregation method in the GCN model. With different information aggregation methods, it can result in different GCN variants. However, no matter which method we choose, the aggregation and transformation layer converts the hidden representation of node *i* in layer *k*, **z**_*i*,*k*_, into the hidden representation in the next layer as **z**_*i*,*k*+1_. We use the output of the last graph convolutional layer, **z**_*i*,*N*_, as the final embedding for that node, **z**_*i*_. Naturally, the input of the first convolutional layer is the original feature vector of each node (Section 2.3). Formally, **z**_*i*,0_ = **x**_*i*_.

### 2.5 Edge prediction from embeddings

In this section, we introduce how to reconstruct edges in the network with the embeddings learned from GCN. We use the bilinear decoder with the following form as the the edge decoder:

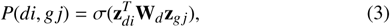

where 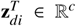 is the learned embedding of a disease node *di*; **z**_*gj*_ ∈ ℝ^*c*^ is the learned embedding of a gene node *gj*; **W**_*d*_ ∈ ℝ^*c*c*^ is the trainable parameter matrix, which models the interaction between each two dimensions of 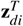 and **z**_*gj*_; σ is the sigmoid function which converts the output value of the edge decoder to the range of (0, 1), as a probability value. This edge decoder is illustrated in Fig. 2 (C). Note that, similar to the graph convolutional neural network model, the parameters of the bilinear decoder model are also shared across different gene-disease pairs, which can effectively reduce the risk of overfitting.

Taking the GCN model and the edge decoder model together, we have the following trainable parameters: (1). The link-type-specific and layer-specific convolutional weight parameters 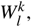, which suggest how to aggregate and transform information from the node’s neighbors. (2). The node-type-specific and layer-specific weight parameters 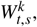, which indicate how to preserve and transform nodes’ self-information from one layer to the next. (3). The weight parameters of the bilinear edge decoder model, **W**_*d*_, which model the interaction between two dimensions of the input embeddings of two nodes. As shown in Fig. 2 (B) and (C), the GCN model and the edge decoder model can be combined together to form an end-to-end model, which takes the raw representation of two nodes and output interaction probability. Consequently, the entire model and all the parameters can be trained in an end-to-end manner.

### 2.6 Model hyper-parameters

In this section, we introduce the hyper-parameters that we chose when building and training the model.

First, we used the cross-entropy loss as the loss function to train the entire model, which has the following form:

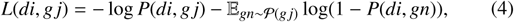

where *di* and *gj* form an edge in the training data. That is, the ground truth value *Y*(*di*, *gj*) = 1 in Fig. 2 (D). By using the cross-entropy loss, we want the model to assign the probabilities for the observed training edges as high as possible while assigning low probabilities for the random edges. Following the previous studies (Trouillon *et al.*, 2016; Zitnik *et al.*, 2018), we used negative sampling to achieve this, which is illustrated by the last term in Eq. (4). For each existing edge (*di*, *gj*), which is a positive sample, we sampled a random edge (*di*, *gn*) by randomly choosing the second node *gn*, which follows the sampling distribution 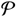. Considering all the edges, we have the final cross-entropy loss of the model as:

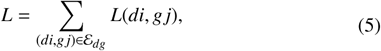

where 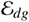 represents all the edges connecting diseases and genes. As we discussed in the previous sections, we trained the model in an end-to-end manner, where the loss function gradient is back-propagated to the parameters in both the GCN model and the edge decoding model. This end-to-end training strategy is more likely to find problem-specific, effective models and embeddings, which has been proved by previous studies (Li *et al.*, 2017; Dai *et al.*, 2017; Umarov *et al.*, 2019; Zou *et al.*, 2019).

In terms of implementation, we set the number of layers as 2, with the dimension of the hidden representation as 64 and the final embedding dimension as 32. We trained the model using Adam optimizer, with the learning rate as 0.001. To reduce overfitting, we used the combination of dropout on the hidden layer unites with the dropout rate as 0.1, and the legendary weight decay method. We initialized the model’s parameters using Xavier initializer. During training, we fed mini-batch of edges to the model, with the batch size as 512. This can reduce the memory requirement and serve as an additional regularizer that further alleviates overfitting. In total, we trained the model for 300 epochs. With the help of a Titan Xp card, we finished the training of a model in 10 hours.

### 2.7 Evaluation criteria

We used the following criteria to evaluate our method and the competing methods: Area Under the Receiver Operating Characteristic curve (AUROC), Area Under the Precision-Recall Curve (AUPRC), Boltzmann-Enhanced Discrimination of ROC (BEDROC), Average Precision at *K* (AP@*K*), and Recall at *K* (R@*K*) score. AUROC is a commonly used criterion in machine learning, which computes the area under the ROC curve. In the disease gene prioritization problem, it can be interpreted as the probability of a true disease-associated gene is ranked higher than a false one selected randomly in a uniform distribution. Similar to AUROC, AUPRC computes the area under the precision-recall curve. BEDROC, proposed to solve the “early recognition” problem, can be interpreted as the probability of a disease-associated gene being ranked higher than a gene selected randomly following a distribution in which top-ranked genes have a higher probability to be chosen. The formal definition of BEDROC can be referred to Truchon and Bayly (2007). P@*K* computes the precision of the prediction if we consider the top *K* predicted associations. Recall at *K* considers the recall score within the top *K* predictions. These five criteria can provide a comprehensive evaluation of the proposed method.

## 3 RESULTS

In this section, we show the performance of the proposed method and the five state-of-the-art methods. We first briefly introduce the five competing methods. Then we introduce the experimental settings in details. After that, we show the performance of all the methods on recovering missing associations, and on discovering associations for novel genes and/or diseases that are not seen in the training. Finally, we demonstrate the effectiveness of the proposed method by investigating the predicted associations on breast cancer.

### 3.1 Compared methods

Five state-of-the-art methods for disease gene prioritization are included in the comparison. The first one is Katz (Singh-Blom *et al.*, 2013), which is a typical network-based method. It computes the node similarity based on the network topology. The similarity matrix is then used to make predictions for disease gene associations. The second one is Catapult (Singh-Blom *et al.*, 2013), another network-based method. It combines the supervised learning with social network analysis, and has been shown to be the state-of-the-art network-based method (Singh-Blom *et al.*, 2013; Natarajan and Dhillon, 2014). This method deploys a biased support vector machine (SVM) as the classifier while the features are derived from random walks in the heterogeneous gene-trait network. It outperformed the previous network-based methods, such as PRINCE and RWRH, significantly. The third one is a very new network-based method, the Graph Convolution-based Association Scoring (GCAS) method (Rao *et al.*, 2018). This method used GCN as a pure network analysis tool which can perform information propagation on the similarity and association networks. Our method differs from GCAS in that we use GCN to integrate information from different sources and learn embeddings specifically for this problem, which are particularly suitable for the downstream edge prediction task. The fourth one is the Inductive Matrix Completion (IMC) method (Natarajan and Dhillon, 2014), which introduced the matrix completion method into the disease gene prioritization field for the first time. It constructs features from genes and diseases from multiple sources, ranging from gene expression array to disease similarity networks. It then learns low-rank latent vectors for diseases and genes which can explain the observed disease-gene associations, taking into consideration features using a linear model. The learned latent vectors are then used for making further predictions. The last one is the very recently developed GeneHound method (Zakeri *et al.*, 2018). It also utilizes the matrix completion method but combines Bayesian approach with matrix completion, which takes the disease-specific and gene-specific information as the prior knowledge. This method has been shown to outperform the legendary Endeavour method significantly (Zakeri *et al.*, 2018).

### 3.2 Experimental settings

We built the dataset from the OMIM database (November 26, 2017). After preprocessing, we constructed a dataset with 12331 genes, 3215 diseases, and 3988 associations. Comprehensive experiments were designed to evaluate the performance of the proposed method. Firstly, we assessed the overall ability to recover the known disease-gene associations using the standard cross-fold validation strategy. During the experiments, we randomly hid 10% associations as the testing set and used the remaining 90% as the training set. This experiment mimics the situation in which partial knowledge about a disease is known (i.e., some associated genes are known) and we want to complete the knowledge by finding out other associated genes. The results are shown in Section 3.3. However, this task is neither the most practically important nor the most challenging one for disease gene prioritization. In reality, researchers are more interested in predicting associations for diseases and/or genes that are not known before. To mimic such situations, we further designed three experiments. The first one is to predict associations for singleton genes (Singh-Blom *et al.*, 2013), which means that the gene has only one associated disease and is not included in the training set (Section 3.4). The second one is to predict associations for new diseases. We excluded all the associations for certain diseases from the training set and challenged different methods to recover these associations (Section 3.5). In the third experiment, we tested the performance of different methods on recovering novel associations, which are defined as the ones that both the disease and the gene are absent in the training set (Section 3.6). Finally, we showed a case study of our predictions for breast cancer in Section 3.7.

### 3.3 Overall performance

We randomly hid 10% associations as the testing set and used the remaining 90% edges as the training data to evaluate the overall performance of different methods on recovering the hidden associations. The performance of different methods is summarized in Table. 1. As shown in the table, the two matrix completion methods, GeneHound and IMC, can outperform the other three network-based methods, GCAS, Catapult and Katz, significantly across different criteria. The main reason is that they can take full advantage of the gene- and disease-specific information while the network-based methods are biased towards the network topology. On the other hand, the proposed method, PGCN, which can utilize both the network topology information and the additional information of the nodes in a systematic and natural way, can outperform all the state-of-the-art methods significantly and consistently across different criteria with a large margin. In terms of AUPRC, PGCN can outperform the second best method by around 10%. We further show the ROC curves and the PRC curves in Fig. 3 (A,B). It is clear that PGCN significantly outperforms all the state-of-the-art methods under all the false positive rates and all the recall values, which suggests an overall much better method. In disease gene prioritization, Recall at *K* is also an important indicator because the top-ranked genes are candidates for further investigation. Fig. 3 (C) shows the recall of different methods when different numbers of top predictions are considered. Interestingly, GCAS can perform quite well when *K* is very small, compared to GeneHound, IMC, Catapult and Katz. Yet PGCN is clearly more sensitive than all the competing methods regardless of the number of top predictions to be considered. All these consistent results demonstrate that the proposed method can outperform the other methods in recovering the hidden associations between diseases and genes.

**Fig. 3:**
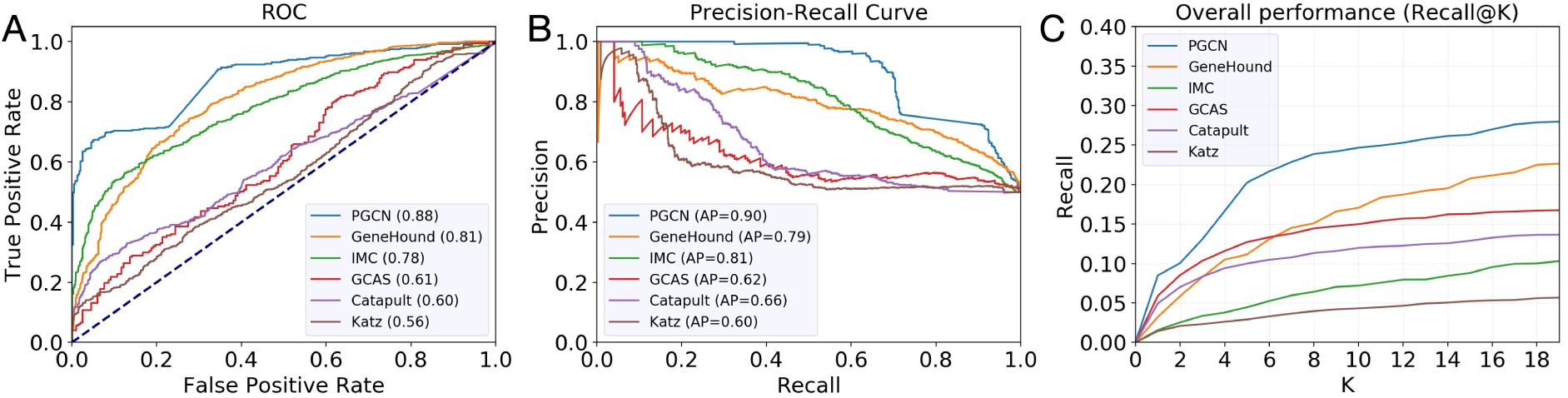
Performance comparison of different methods. For drawing the ROC curves (A) and PRC curves (B), we randomly selected the same number of negative samples to the positive samples from the network. As shown in the figure, the proposed method, PGCN, can outperform the competing methods significantly in terms of AUROC and AUPRC. (C) The performance of different methods in terms of recall at *K*. This criterion suggests the probability of an actual association being retrieved when checking the top-*K* predictions. The proposed method, PGCN, can outperform all the other compared methods regardless of the value of *K*.

**Table 1.** The overall performance of the five compared methods. Under each criterion, the method with the best performance is in bold and the second best is underlined.

| Method | AUROC | AUPRC | AP@200 | BEDROC |
| --- | --- | --- | --- | --- |
| PGCN | **0.877** | **0.896** | **0.976** | **0.987** |
| GeneHound | <u>0.805</u> | 0.793 | 0.831 | 0.908 |
| IMC | 0.780 | <u>0.809</u> | <u>0.928</u> | <u>0.965</u> |
| GCAS | 0.614 | 0.623 | 0.753 | 0.813 |
| Catapult | 0.597 | 0.657 | 0.783 | 0.884 |
| Katz | 0.557 | 0.596 | 0.595 | 0.790 |

### 3.4 Performance on singleton genes

Following the idea of Singh-Blom *et al.* (2013), we checked the performance of different methods on predicting the associations of singleton genes, which are defined as those genes with only one link in the database. In our experiment, the only links for the singleton genes were removed from training, which means that the methods needed to predict the associations “from scratch”. We used recall at *K* to evaluate different methods, which is a difficult measurement because each test gene has one and only one true association. As shown in Fig. 4 (A), PGCN consistently recovers the missing associations for singleton genes better than other methods. We also noticed that the network information is very important when *K* is small (between 1 and 10) because the improvement of PGCN over the network-based method (e.g., Katz) is not large, which is consistent with the previous findings (Natarajan and Dhillon, 2014). However, as the number of top predictions being considered increases, the disease- and gene-specific information plays more and more important role, which leads to significantly better recall when *K* is large.

**Fig. 4:**
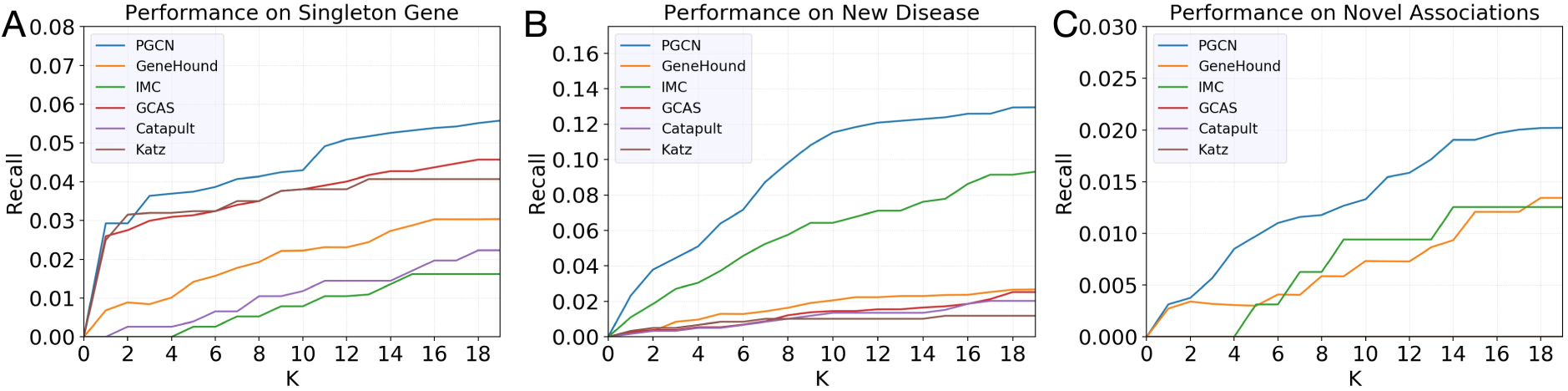
Performance in terms of recall at *K* of different methods on recovering associations for new genes and/or diseases. (A) Performance comparison of different methods on the singleton genes association prediction. (B) Performance comparison of different methods on the new disease association prediction. (C) Performance comparison of different methods on the novel association prediction.

### 3.5 Performance on new diseases

Next, we evaluated the ability of different methods on predicting associations for novel diseases for which no associated genes are known. For a novel disease, all of its associations with genes were removed during training and different methods were challenged to recover those missing associations. This task is considerably less difficult in terms of recall than recovering the associations for singleton genes because a disease can be associated with more than one genes. At the same time, this task is practically important because it is directly related to the molecular diagnosis for human diseases. As shown in Fig. 4 (B), IMC can outperform all the other previous methods with a large margin. The reason is that IMC is based on matrix completion techniques, which can effectively incorporate the disease-specific information (Natarajan and Dhillon, 2014). Our method, however, can not only incorporate disease- and gene-specific information, but also the known disease-gene associations in a unified framework. Furthermore, our method trains the disease and gene embeddings and link prediction in an end-to-end manner, and thus further significantly improves the performance over IMC.

To further understand how our method works, we investigate a disease, atrioventricular septal defect-4 (AVSD4), for which we removed its only associated gene, *GATA4*, during training and PGCN successfully recovered it with the highest score. The link between AVSD4 and *GATA4* is built through another disease, ventricular septal defect-1 (VSD1), which is known to be associated with *GATA4*. Our method detected the similarity between the two diseases, AVSD4 and VSD1, according to their embeddings learned by our method, which is illustrated in Fig. 6 (B). However, this similarity is very difficult to be detected because in the disease similarity network, the two diseases have a wrong similarity score of 0, which suggests that they are two completely irrelevant diseases. Therefore, all the network-based methods failed to predict the association between AVSD4 and *GATA4*. Our method, on the contrary, systematically incorporates not only the network topology, but also the disease-specific information. In this particular case, the disease-specific information plays an important role in the disease embedding and thus PGCN was able to detect the similarity between the two diseases in the embedding space, which led to the correct prediction on the association between AVSD4 and *GATA4*.

**Fig. 5:**
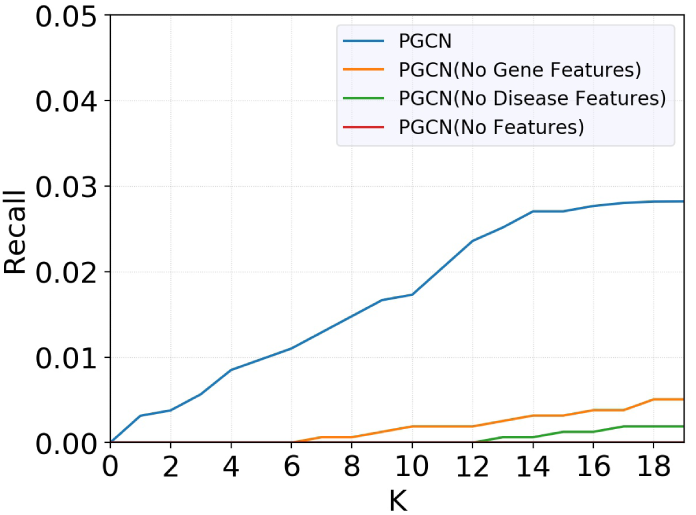
The importance of the disease- and gene-specific information. We show the performance of the proposed method on the novel association prediction when eliminating the feature vectors for different types of nodes. “PGCN” is the proposed method, which trains the model with both the disease features and gene features. “PGCN (No Gene Features)” trains the model without the features for the gene nodes. “PGCN (No Disease Features)” trains the model without the features for the disease nodes. “PGCN (No Features)” only uses the network topology information, without using any features for disease or gene nodes.

**Fig. 6:**
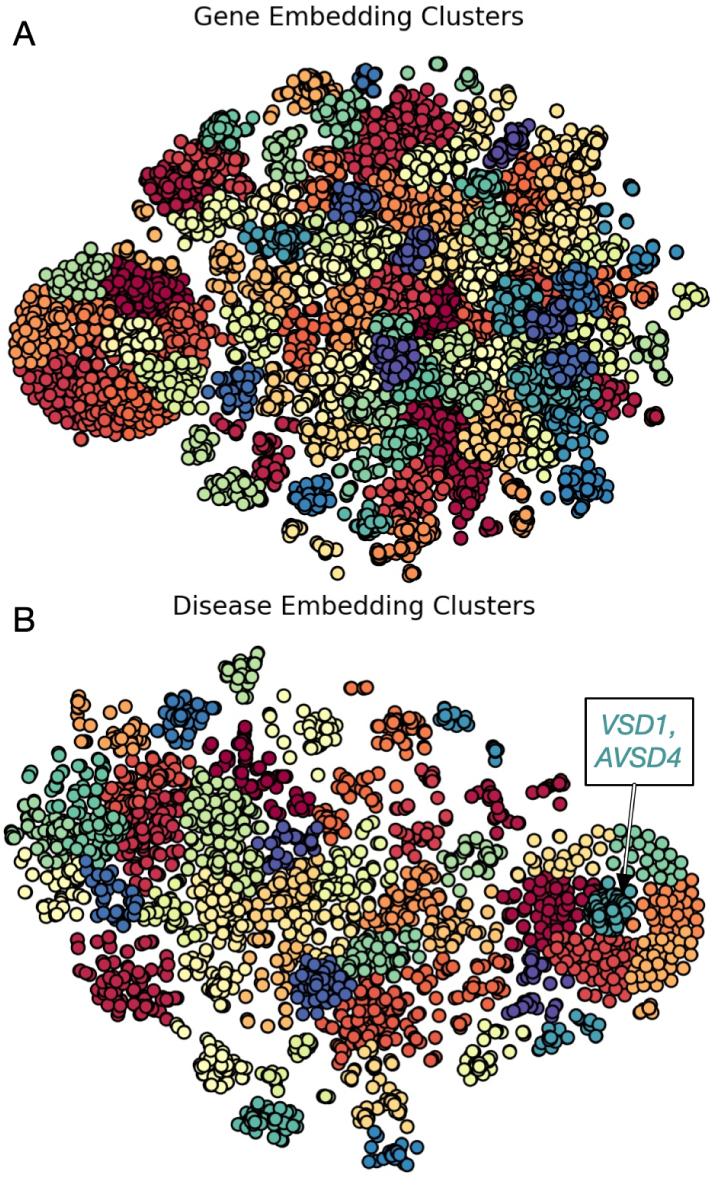
Visualization of the clustering of embeddings in 2D space using t-SNE.

### 3.6 Performance on novel associations

We then evaluated the prediction performance of different methods for novel associations, which are defined to be the associations between a disease and a gene, both of which have no association in the training set. This is the most stringent and challenging requirement. In order for a method to recover such associations, neither the disease end nor the gene end of the association can be directly used. The method must be powerful enough to effectively use the disease- and gene-specific information, and propagate the information through other diseases, genes, and their associations in the heterogeneous network. The results are shown in Fig. 4 (C). As expected, the recall values of all the methods have a clear drop comparing to the two previous tasks. We found that the three network-based methods did not perform well in this task as they were unable to recall any true associations. We suspect that the main reason for this is that the definition of novel associations makes network propagation alone extremely difficult. To support this view, the two matrix completion methods, which can take advantage of the specific information of genes and diseases, performed much better than the network-based methods. Our method consistently outperforms all the competing methods, and the improvement increases with a larger *K*.

### 3.7 Case study

As a case study, we investigated the top 10 associations for breast cancer. Among these 10 genes, other than the four ground-truth breast cancer-related genes reported in the OMIM dataset, our model also predicted three interesting genes: *Axin2*, *TLR4*, and *PTPRJ*, which were reported to be related to breast cancer. For example, *Axin2* was found to be included in the *Wnt*/β*-catenin*/*Axin2* pathway, which can regulate the breast cancer invasion and metastasis (Li *et al.*, 2016); *TLR4* was found to be overexpressed in the majority of the breast cancer samples and also related to the metastasis of breast cancer (Volk-Draper *et al.*, 2014); and *PTPRJ* forms *DEP-1*/*PTPRJ*/*CD148*, which is receptor-like protein tyrosine phosphatases (PTP), that was found to be mutated or deleted in human breast cancer (Spring *et al.*, 2015). These results suggest the potential application of our method on discovering new genes related to complex human diseases.

## 4 DISCUSSION

### 4.1 Importance of disease- and gene-specific information

To expand the analysis for the importance of the disease- and gene-specific information, we further investigated its contributions to the prediction performance of our method. Focusing on the novel association prediction task, we excluded the disease features, the gene features, and both of them, respectively, and evaluated the performance of the corresponding models. As shown in Fig. 5, both the disease features and the gene features are very important for the proposed method. If we exclude either one of them, the performance will degrade significantly. If we exclude both of them, the model cannot recall anything when *K* is in the range of (1, 19). On the other hand, disease features are more important than the gene features as the model with the disease features begins to recall some true associations when *K* = 7 while the model with the gene features begins to recall some true associations when *K* = 13. The reason may be that the gene network we used is HumanNet, which is a very informative database that was built from multiple data sources.

### 4.2 Biological meaning of embeddings

To gain insights into how the final embeddings represent the gene and the disease features, we mapped the 32-dimensional vector of each node into a 2-dimensional space using t-SNE (Maaten and Hinton, 2008) for visualization (Fig. 6). From these 2-dimensional data, we observed that many points are located closely to some other points and they form clusters of a wide range of sizes in both the gene feature space and the disease feature space. Since closely located data points suggest that the corresponding features be biologically similar in our embedding, we analyze the extent to which closely located data points in the low dimensional space represent the biological association of the corresponding features.

To this end, we first clustered the data points into 100 groups for the gene node embedding and 50 groups for the disease node embedding using hierarchical clustering. To analyze the functional association for gene features, we mapped genes in each cluster to biological pathways that they are associated with using the KEGG pathway data (Kanehisa and Goto, 2000) and evaluated their statistical significance. We found that all of the 100 clusters have statistically significant levels of association with biological function (*p* < 0.05; hypergeometric test). Notably, the cluster which includes RPL3L over-represents the genes involved in the formation of the ribosome (*p* < 10^−82^), while the one including H2AFX has a disproportionately large number of genes involved in the DNA repair response of systemic lupus erythematosus (*p* < 10^−39^).

For the analysis of the disease node embedding, we used the Human Phenotype Ontology (HPO) dataset (Köhler *et al.*, 2019) to associate each disease with corresponding HPO phenotypic abnormality terms. Similar to the gene-function association analysis, all of the 50 clusters are found to have statistically significant number of diseases with association to some sort of phenotypic abnormalities (*p* < 0.01). In particular, we found that the cluster including Parkinson Disease, Late-onset (OMIM:168600) is enriched in diseases that are associated with slow movements (*p* < 10^−22^), while the cluster with Neuropathy, Hereditary Sensory And Autonomic, Type Ii (OMIM:201300) over-represents genes associated with muscular hypotonia (*p* < 10^−10^).

These results indicate the ability of our method to generate embeddings that preserve the gene and the disease associations that are critical to the disease gene prioritization task. They also highlight the possibility to interpret the gene node and the disease node embeddings in a biologically meaningful way, which is essential to gain biomedical insights into novel disease-gene association.

## 5 CONCLUSION

In this paper, we proposed a novel, unified framework for disease gene prioritization. Our method automatically learns the embedding of diseases and genes by systematically incorporating the topology of the heterogeneous network, the neighborhood of the diseases and genes, and the disease- and gene-specific information. The embeddings and the association prediction models are trained in an end-to-end manner. Extensive experiments demonstrate the power of our method on recovering missing associations, and on discovering associations for novel genes and/or diseases that are not seen in the training. Our framework is generic and can be readily applied to tackle other important problems in computational biology, such as drug disease association (Pushpakom *et al.*, 2019) and homolog detection for protein structure prediction (Cui *et al.*, 2016).

## ACKNOWLEDGMENTS

This work was supported by the King Abdullah University of Science and Technology (KAUST) Office of Sponsored Research (OSR) under Awards No. FCC/1/1976-17-01, FCC/1/1976-18-01, FCC/1/1976-23-01, FCC/1/1976-25-01, FCC/1/1976-26-01, and URF/1/3412-01.

